# Post-mortem molecular profiling of three psychiatric disorders

**DOI:** 10.1101/061416

**Authors:** Ryne C. Ramaker, Kevin M. Bowling, Brittany N. Lasseigne, Megan H. Hagenauer, Andrew A. Hardigan, Nick S. Davis, Jason Gertz, Preston M. Cartagena, David M. Walsh, Marquis P. Vawter, Edward G. Jones, Alan F. Schatzberg, Jack D. Barchas, Stan J. Watson, Blynn G. Bunney, Huda Akil, William E. Bunney, Jun Z. Li, Sara J. Cooper, Richard M. Myers

## Abstract

**Background:** Psychiatric disorders are multigenic diseases with complex etiology contributing significantly to human morbidity and mortality. Although clinically distinct, several disorders share many symptoms suggesting common underlying molecular changes exist that may implicate important regulators of pathogenesis and new therapeutic targets.

**Results:** We compared molecular signatures across brain regions and disorders in the transcriptomes of postmortem human brain samples. We performed RNA sequencing on tissue from the anterior cingulate cortex, dorsolateral prefrontal cortex, and nucleus accumbens from three groups of 24 patients each diagnosed with schizophrenia, bipolar disorder, or major depressive disorder, and from 24 control subjects, and validated the results in an independent cohort. The most significant disease differences were in the anterior cingulate cortex of schizophrenia samples compared to controls. Transcriptional changes were assessed in an independent cohort, revealing the transcription factor *EGR1* as significantly down regulated in both cohorts and as a potential regulator of broader transcription changes observed in schizophrenia patients. Additionally, broad down regulation of genes specific to neurons and concordant up regulation of genes specific to astrocytes was observed in SZ and BPD patients relative to controls. We also assessed the biochemical consequences of gene expression changes with untargeted metabolomic profiling and identified disruption of GABA levels in schizophrenia patients.

**Conclusions:** We provide a comprehensive post-mortem transcriptome profile of three psychiatric disorders across three brain regions. We highlight a high-confidence set of independently validated genes differentially expressed between schizophrenia and control patients in the anterior cingulate cortex and integrate transcriptional changes with untargeted metabolite profiling.

## Background

Schizophrenia (SZ), bipolar disorder (BPD), and major depressive disorder (MDD) are multigenic diseases with complex etiology and are large sources of morbidity and mortality in the population. All three disorders are associated with high rates of suicide, with ∼90% of the ∼41,000 people who commit suicide each year in the U.S. having a diagnosable psychiatric disorder [1]. Notably, while clinically distinct, these disorders also share many symptoms, including psychosis, suicidal ideation, sleep disturbances and cognitive deficits [2–4]. This phenotypic overlap suggests potential common genetic etiology, which is supported by recent large-scale genome-wide association studies [5–8]. However, this overlap has not been fully characterized with functional genomic approaches. Current therapies for these psychiatric disorders are ineffective in many patients and often only treat a subset of an individual patient’s symptoms [9]. Approaches targeting the underlying molecular pathologies within and across these types of disorders are necessary to address the immense burden of psychiatric disease around the world and improve care for the millions of people diagnosed with these conditions.

Previous studies [10–14] analyzed brain tissue with RNA sequencing (RNA-seq) in SZ and BPD, and identified altered expression of GABA-related genes in the superior temporal gyrus and hippocampus, as well as differentially expressed genes related to neuroplasticity and mammalian circadian rhythms. Our study focused on the anterior cingulate cortex (AnCg), dorsolateral prefrontal cortex (DLPFC), and nucleus accumbens (nAcc), regions which are often associated with mood alterations, cognition, impulse control, motivation, reward, and pleasure – all behaviors known to be altered in psychiatric disorders [15,16]. To assess gene expression changes associated with psychiatric disease in these three brain regions, we performed RNA-seq on macro-dissected post-mortem tissues in four well-documented cohorts of 24 patients each with SZ, BPD, MDD and 24 controls (CTL) (96 individuals total). Additionally, we conducted metabolomic profiling of AnCg tissue from the same subjects. RNA-seq analysis revealed common expression profiles in SZ and BPD patients supporting the notion that these disorders share a common molecular signature. Transcriptional changes were most pronounced in the AnCg with SZ and BPD exhibiting strongly correlated differences from CTL samples. Differentially expressed genes were associated with cell-type composition with BPD and SZ samples showing decreased expression of neuron-specific genes. We validated this result with RNA-seq data from an independent cohort of 35 cases each of SZ, BPD, and CTL post-mortem cingulate cortex samples from the Stanley Neuropathology Consortium Integrative Database (SNCID; http://sncid.stanleyresearch.org) Array Collection. We present a set of validated genes differentially expressed between SZ and CTL patients, perform an integrated analysis of metabolic pathway disruptions, and highlight a role for the transcription factor, *EGR1*, whose down-regulation in SZ patients may drive a large portion of observed transcription changes.

## Methods

See Supplemental Methods for additional detail.

### Patient Sample Collection and Preparation

Sample collection, including human subject recruitment and characterization, tissue dissection, and RNA extraction, was described previously [17,18] as part of the Brain Donor Program at the University of California, Irvine, Department of Psychiatry and Human Behavior (Pritzker Neuropsychiatric Disorders Research Consortium) under IRB approval. In brief, coronal slices of the brain were rapidly frozen on aluminum plates that were previously frozen to −120°C and dissected as described previously [19]. All samples were diagnosed by psychological autopsy, which included collection and analyses of medical and psychiatric records, toxicology, medical examiners’ reports, and 141-item family interviews. Agonal state scores were assigned based on a previously published scale [20]. Controls were selected based upon absence of severe psychiatric disturbance and mental illness within first-degree relatives.

We obtained fastq files from RNA-seq experiments for our validation cohort from the Stanley Neuropathology Consortium Integrative Database (SNCID; http://sncid.stanleyresearch.org) Array Collection comprising 35 cases each of SZ, BPD, and CTL of post-mortem cingulate cortex with permission on June 30, 2015. For our analysis, we included the 27 SZ, 26 CTL, and 25 BPD SNCID samples that were successfully downloaded and represented unique samples. SNCID RNA-seq methodology and data processing are described in detail in a previous publication that makes use of the data [10].

### RNA-seq and Data Processing

To extract nucleic acid, 20 mg of post-mortem brain tissue was homogenized in Qiagen RLT buffer + 1% BME using an MP FastPrep-24 and Lysing Matrix D beads for three rounds of 45 seconds at 6.5 m/s (FastPrep homogenizer, lysing matrix D, MP Bio). Total RNA was isolated from 350 μL tissue homogenate using the Norgen Animal Tissue RNA Purification Kit (Norgen Biotek Corporation). We made RNA-seq libraries from 250 ng total RNA using polyA selection (Dynabeads mRNA DIRECT kit, Life Technologies) and transposase-based non-stranded library construction (Tn-RNA-seq) as described previously [21]. To mitigate potentially confounding batch affects in sample preparation we randomly assigned samples from all brain regions and disorders into batches of 24 samples. We used KAPA to quantitate the library concentrations and pooled 4 samples in order to achieve equal concentration of the four libraries in each lane. Pools were determined by random from the 291 samples. Samples were also randomly selected for pooling in an effort to limit potentially confounding sequencing batch effects. The pooled libraries were sequenced on an Illumina HiSeq 2000 sequencing machine using paired-end 50 bp reads and a 6 bp index read, resulting in an average of 48.2 million reads per library. To quantify the expression of each gene in both Pritzker and SNCID datasets, RNA-seq reads were processed with aRNApipe v1.1 using default settings [22]. Briefly, reads were aligned and counted with STAR v2.4.2a to all genes annotated in GRCh37_E75 [23]. All alignment quality metrics were obtained from the picard tools module (http://broadinstitute.github.io/picard/) available in aRNApipe. Genes expressed from the X and Y chromosomes were omitted from the study.

Quantitative PCR (qPCR) was performed on 10 SZ and 10 CTL patients to validate *EGR1* RNA-seq measurements. RNA was extracted as described above from tissue lysates a second time. Reverse transcription was performed on 250 ng of input RNA with the Applied Biosystems high capacity cDNA reverse transcription kit. Validated Taqman assays for EGR1 (Hs00152928_m1) and the housekeeper genes GAPDH (Hs02758991_g1) and ACTB (Hs01060665_g1) were used for qPCR. cDNA was diluted by a factor of 10 before use as input for the Taqman assay. The qPCR reaction was performed on an Applied Biosystems Quant Studio 6 Flex system using the recommended amplification protocol for Taqman assays.

### Sequencing Data Analysis

All data analysis in R was performed with version 3.1.2.

#### Differential Expression Analysis and Normalization

To examine gene expression changes, we employed the R package DESeq2 [24] (version 1.6.3), using default settings, but employing likelihood ratio test (LRT) hypothesis testing, and removing non-convergent genes from subsequent analysis. Genes differentially expressed between each disorder and CTL samples, by brain region, were identified with DESeq2 (adjusted p-value<0.05), including age, brain pH, PMI, and percentage of reads uniquely aligned (PRUA) as covariates (Full Model: ∼Age+PMI+pH+PRUA+Disorder, Reduced Model: ∼ Age+PMI+pH+PRUA). For downstream heatmap visualization, PCA, and cell-type analysis, genes underwent a log-like normalization using DESeq2’s varianceStabilizingTransformation function and were corrected for PRUA by computing residuals to a linear model regressing PRUA on normalized gene expression level with the R lm function unless otherwise specified. DESeq2’s default independent filtering method was used to remove genes with an insufficient expression level from further analysis.

#### PCA and Hierarchical Clustering

PCA analysis was performed in R on normalized data using the prcomp() command. Hierarchical clustering of normalized gene expression data was done in R with the hclust command (method=“ward”, distance=“Euclidean”)

#### Pathway Enrichment Analysis

Pathway analysis was conducted using the web-based tool LRPath [25] using all GO term annotations, adjusting to gene read count with RNA-Enrich, including directionality and limiting maximum GO term size to 500 genes. GO term visualization was performed using the Cytoscape Enrichment Map plug-in [26]. The Genesetfile (.gmt) GO annotations from February 1, 2017 were downloaded from http://download.baderlab.org/EM_Genesets/. The LRPath output was parsed and used as an enrichment file with all upregulated pathways colored red and all downregulated pathways colored blue, regardless of degree of upregulation. Mapping parameters were; p-value cutoff = 0.005, FDR cutoff = 0.1 and Jaccard coefficient > 0.3. Resulting networks were exported as PDFs. Summary terms were added to the plot based on the GO terms in those clusters. In order to assess overlap between significant GO terms in our analysis and the GWAS study described by the Psychiatric Genomics Consortium, we downloaded the p-values reported for Schizophrenia hits from their Supplemental Table 4, which contained 424 significant GO terms. We used a chi-squared test to assess significant overlap between the two groups. Our Supplemental Table 5 reports the p-values measured in SZ based on this study along with those calculated in our analysis.

#### EGR1 ChIP-seq peak analysis

Narrow peak bed files filtered to optimal IDR peaks were obtained from the ENCODE data portal (www.encodeproject.org) for *EGR1* ChIP-seq data in GM12878, H1-hESC, and K562 cell lines (ENCODE file IDs: ENCFF002CIV, ENCFF002CGW, ENCFF002CLV). Consensus *EGR1* peaks were identified by intersecting peaks from all three cell lines, which resulted in a final list of 4,121 peaks common to all cell lines (minimum overlap of 1 bp). The distance from each annotated transcription start site (TSS) to the nearest consensus *EGR1* peak was computed based on TSSs annotated in the ENSEMBL gene transfer format (GTF) file from the Ensembl data release 75 (GRCh37_E75).

#### Cell-Specific Enrichment Analysis

Sets of genes uniquely expressed by several brain cell-types were obtained from figure 1B in Darmanis et. al [27]. An index for each cell-type was created by calculating the median normalized expression value for each set of cell-type associated genes. Index values were compared across patient clusters by non-parametric rank sum tests and spearman correlation with top principal components. To validate our method, we calculated cell-type specific indices from an independent cohort of previously published purified brain cells [28,29]. FPKM-normalized gene expression data was obtained from supplemental table 4 of Zhang et. al. (2014) and cell-type indexes were calculated as described above. To examine index performance in mixed cell populations, we obtained fastq files for neuron and astrocyte-purified brain samples from GEO accession GSE73721 and generated raw count files as described above. We next mixed expression profiles *in silico* by performing random down-sampling of neuron and astrocyte count levels and summing the results such that mixed populations containing specific proportions of counts from neuron- and astrocyte-purified tissue were generated. For example, to generate an 80/20 neuron to astrocyte mixture, neuron and astrocyte count columns (which started at an equivalent number of 5,759,178 aligned reads) were randomly down-sampled to 4,607,342 and 1,151,836 counts respectively and summed across each gene to result in a proportionately mixed population of aligned count data simulating heterogeneous tissue. Then we calculated a neuron/astrocyte index ratio capable of predicting the *in silico* mixing weights. Briefly, we assumed index values for mixed cell populations were directly proportional to mixing weights of their respective purified tissue, thus the predicted cell proportion for a given cell type was simply calculated as: predicted cell proportion=observed index value/purified tissue index value

**Figure 1.**
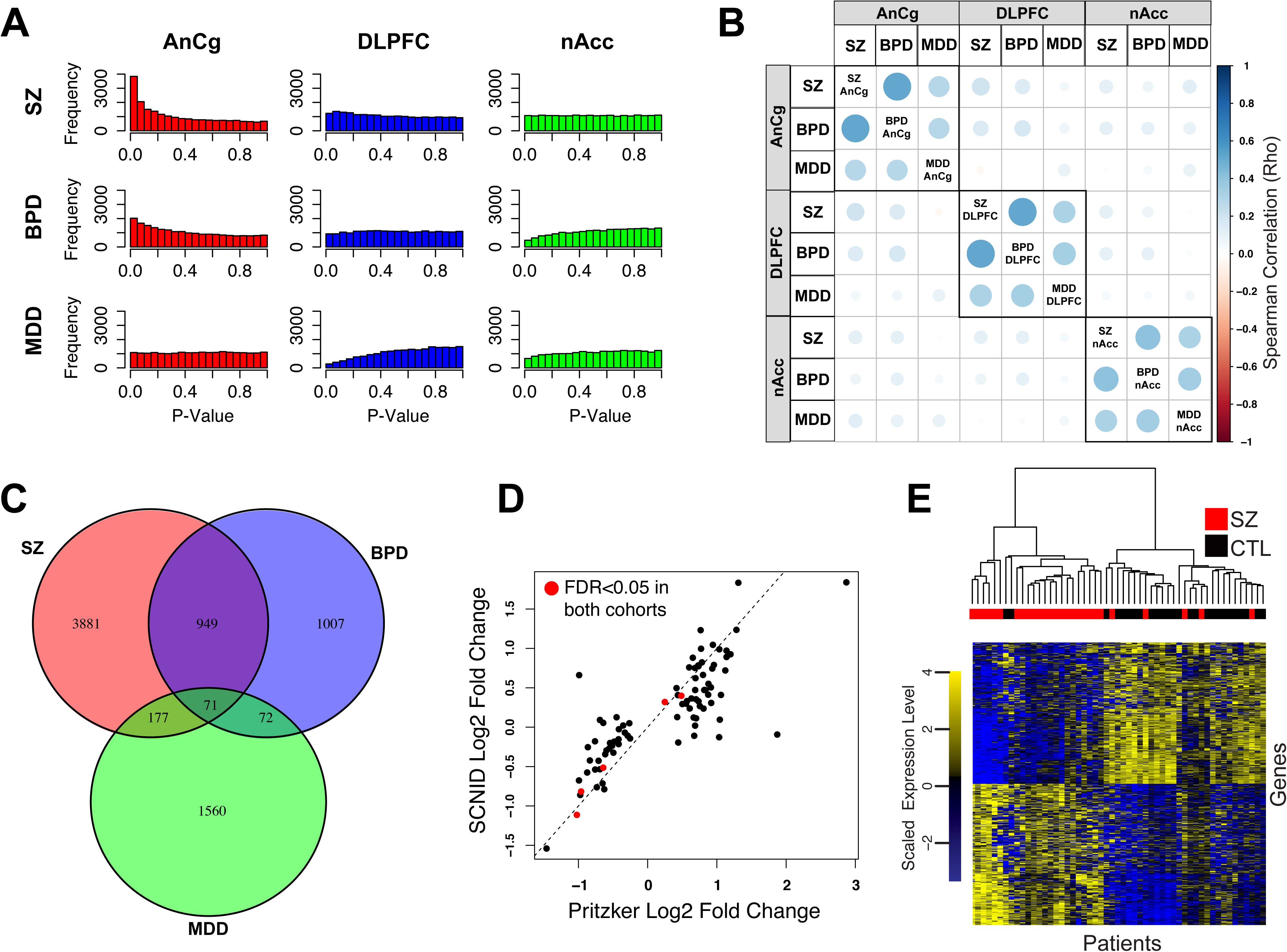
(A) Histograms of case vs. control differential expression (DESeq2 p-values) for SZ (red), BPD (blue), and MDD (green) in each brain region assayed. A minimum DESeq2 base mean of 10 was required for inclusion. (B) Pairwise spearman correlations of log_2_ fold gene expression changes between each disorder and CTL in each brain region. Circle sizes are scaled to reflect absolute Spearman correlations. (C) Venn diagram showing overlap of genes differentially expressed between SZ (red), BPD (blue), MDD (green) vs. CTL at p-value<0.05 in the AnCg. (D) Log2 fold expression change correlation of 87 genes with FDR<0.05 comparing SZ and CTL (AnCg) in the Pritzker dataset with the SNCID dataset (Spearman coefficient=0.812, p-value<0.0001). Genes differentially expressed at an FDR<0.05 in both cohorts are identified with red circles. (E) Hierarchical clustering 27 SZ and 26 CTL tissues in the SNCID dataset using variance-stabilized expression of 1003 genes differentially expressed between SZ and CTL in the AnCg (uncorrected p-value<0.05) in the Pritzker dataset. CTL (black), SZ (red), lowly expressed genes (blue pixels), highly expressed genes (yellow pixels).

To insure cell-type predictive power was unique to indices derived from Darmanis et. al genes, we generated indices from 10,000 randomly sampled gene sets of equivalent size and examined their performance in predicting *in silico* mixing weights. Mean squared prediction errors (MSE) were calculated for each of the 10,000 null indices and compared to the MSE of Darmanis et. al.-derived indices.

Cell type deconvolution analysis was confirmed using a previously published algorithm implemented in the R package deconRNAseq [30]. The “datasets” input to the deconRNAseq function was a normalized count matrix of all AnCg brain samples and the “signatures” input consisted of a normalized count matrix of astrocyte, neuron, microglia, and oligodendrocyte dissected cells from the GEO accession GSE73721 previously described.

Enrichment analysis for extreme fold change was performed by randomly sampling the fold changes of 1000 null gene sets equivalent in size and expression level (allowing 5% error) to the neuron and astrocyte specific gene sets. The median fold change of each 1000 null gene set was compared to the observed median fold change for neuron and astrocyte gene sets respectively.

### Metabolomics

#### Sample preparation

Sections of approximately 100 mg of frozen tissue were weighed and homogenized for 45 seconds at 6.5M/s with ceramic beads in 1 mL of 50% methanol using the MP FastPrep-24 homogenizer (MP Biomedicals). A sample volume equivalent to 10 mg of initial tissue weight was dried down at 55°C for 60 minutes using a vacuum concentrator system (Labconco). Derivatization by methoximation and trimethylsilylation was done as previously described [31].

We analyzed technical replicates of each tissue sample, in randomized order.

#### GCxGC-TOFMS analysis

All derivatized samples were analyzed on a Leco Pegasus 4D system (GCxGC-TOFMS), controlled by the ChromaTof software (Leco, St. Joseph, MI). Samples were analyzed as described previously [31] with minor modifications in temperature ramp.

#### Data analysis and metabolite identification

Peak calling, deconvolution and library spectral matching were done using ChromaTOF 4.5 software. Peaks were identified by spectral match using the NIST, GOLM [32], and Fiehn libraries (Leco), and confirmed by running derivatized standards (Sigma). We used Guineu for multiple sample alignment [33].

#### Integrated Pathway Analysis

Altered metabolites and genes were analyzed for enrichment in KEGG pathways containing both metabolite and gene features. A non-parametric, threshold free pathway analysis similar to that of a previously described method [34] was first performed on metabolite and gene expression data separately. Our method builds on the principle described by Subramanian that implements a one-tailed Wilcox test to identify pathways enriched for low p-values. Instead of just accounting for enrichment at the gene level, we use metabolite or gene p-value ranks within each pathway compared to remaining non-pathway metabolites or genes with a one-tailed Wilcox test to test the hypothesis that elements of a given pathway may be enriched for lower p-value ranks than background elements. Metabolite and gene p-values were subsequently combined to provide an integrated enrichment significance p-value using Fisher’s method. Pathways had to contain greater than 5 genes and 1 metabolite measured in our dataset to be included in the analysis. Table 10 lists p-values for enriched pathways based on genes, metabolites or combined.

## Results

### Region-specific gene expression in control and psychiatric brain tissue

We collected post-mortem human brain tissue, associated clinical data, including age, sex, brain pH, and post-mortem interval (PMI), and cytotoxicology results (Tables S1-2) for matched cohorts of 24 patients each diagnosed with SZ, BPD, or MDD, as well as 24 control individuals with no personal history of, or first-degree relatives diagnosed with, psychiatric disorders. Importantly, to limit the effect of acute patient stress at the time of death as a potential confounder we included only patients with an agonal factor score of zero and a minimum brain pH of 6.5 [18]. Using RNA-seq [21], we profiled gene expression in three macro-dissected brain regions (AnCg, DLPFC, nAcc). After quality control, we analyzed 57,905 ENSEMBL genes in a total of 281 brain samples (Table S3).

To examine heterogeneity across brain regions and subjects, we performed a principal component analysis (PCA; Figure S1A) of all genes. The first principal component (PC1, 21.8% of the variation) separates cortical AnCg and DLPFC samples from subcortical nAcc samples. Examination of the first and second principal components for disorder associations reveals a separation of some SZ and BPD samples from all other samples (Figures S1B and S2A-C). However, in agreement with previously reported post-mortem brain RNA sequencing studies [14], we found several principal components to be highly correlated with quality metrics including the percentage of reads uniquely aligned and percentage of reads aligned to mitochondrial sequence (absolute Rho>0.5, FDR<1E-16, Table S4). To reduce the potentially confounding effects of sample quality, we repeated the PCA on expression data normalized to the percentage of reads uniquely aligned for each sample and found that global disease-specific expression differences were significantly reduced and PC1 primarily separated nAcc samples from AnCg and DLPFC brain regions (Figures S1C and S2D-I).

### Disease-specific gene expression in control and psychiatric brains

We next applied DESeq2 [24], a method for analyzing differential sequence read count data, to identify genes differentially expressed across disorders within each brain region after correcting for biological and technical covariates. The largest number of significant expression changes occurred in AnCg between SZ and CTL individuals (87 genes, FDR<0.05, Figure 1A). Pathway enrichment analysis of differentially expressed genes between SZ and CTL patients revealed 935 gene ontology (GO) terms with an FDR<0.05 (Table S5) (122 GOCC, 159 GOMF, and. 654 GOBP). Significant GO terms fall into the broad categories of synaptic function and signaling (e.g. neurotransmitter transport, ion transport, calcium signaling) (Figure S3). These terms overlap significantly with those identified by the Psychiatric Genomics Consortium in their analysis of GWAS implicated genes [35] with 68 GO terms meeting a p-value cutoff of <0.05 in both datasets (p-value<0.0001, Chi-square test). Additionally, nine genes were differentially expressed between SZ and CTL individuals in DLPFC. Three of these were also identified in AnCg: *SST*, *PDPK2P* and *KLHL14*. No genes had an FDR<0.05 when comparing BPD or MDD samples to CTLs in any brain region, or comparing SZ and CTL tissues in nAcc (Table S6). To examine potential common gene expression patterns between the psychiatric disorders, we performed pair-wise correlation calculations of all gene log2 fold changes for each disorder versus controls in each brain region. Of the nine case-control comparisons (for three regions and three diseases), a particularly strong correlation is observed between BPD and SZ compared to either SZ or BPD and MDD in each brain region (Figure 1B). In the AnCg, BPD and SZ share 1,020 common genes differentially expressed at an uncorrected DESeq2 p-value<0.05 compared to only 248 and 143 genes shared between MDD and SZ or BPD respectively (Figure 1C). This strong overlap between BPD and SZ (Fisher’s exact p-value<1E-16) indicates that although expression changes are weaker in BPD they follow a trend similar to those identified in SZ.

Because previous post-mortem analyses have been limited by, and are particularly vulnerable to, biases inherent to examining a single patient cohort, we sought to generate a robust set of SZ associated genes by validating our observed expression changes in an independent cohort. To accomplish this, we examined gene expression differences in the AnCg between SZ and CTL samples in the SNCID RNA-seq Array dataset [13], revealing 1,003 genes altered (DESeq2 uncorrected p-value<0.05) in both datasets (Fisher’s p-value<1E-16, Table S7). The magnitude and direction of change in significant genes in the Pritzker dataset were highly correlated with the SNCID dataset (Rho=0.202, p-value<1E-16), particularly in 87 genes that met a cutoff of FDR<0.05 (Rho=0.812, p-value<1E-16; Figure 1D). We performed hierarchical clustering of SZ and CTL samples in the SNCID validation cohort using the 1,003 genes differentially expressed, at the less stringent threshold, p-value<0.05, between SZ and CTL in the Pritzker dataset (Figure 1E), and found these genes successfully distinguished the two disease groups with only 5 out of 27 SZ and 2 out of 26 CTL samples misclassified.

Of particular interest are the 5 genes significant at a FDR<0.05 in both cohorts including a nearly 2-fold decrease in expression of the transcription factor *EGR1* (Table S7A, Figure 2A). Quantitative PCR (qPCR) validation confirmed reduced *EGR1* expression in SZ samples (Figure 2B, Wilcox p-value=4.33×10E-5). *EGR1*, a zinc finger transcription factor, has been recently implicated in SZ by a GWAS study [5], thus we sought to investigate whether loss of *EGR1* expression might be associated with transcriptional changes observed in the AnCg of SZ patients using publicly available genome-wide occupancy data from the ENCODE consortium (https://www.encodeproject.org). To obtain high confidence *EGR1* binding sites we intersected chromatin immunoprecipitation sequencing (ChIP-Seq) peaks derived from the H1-hESC, K562, and GM12878 cell lines. We found that genes with a transcription start site (TSS) within 1 kb of an *EGR1* binding site had significantly lower DESeq2 p-values (Wilcox p-value=9.68E-5) and reduced expression in SZ versus CTL (Wilcox p-value=7.69E-15) compared to genes whose TSSs were greater than 1 kb from an *EGR1* binding site. A monotonic decrease in this effect was observed as the distance threshold used for this comparison was increased from 1 kb to 50 kb (Figure 2C).

**Figure 2.**
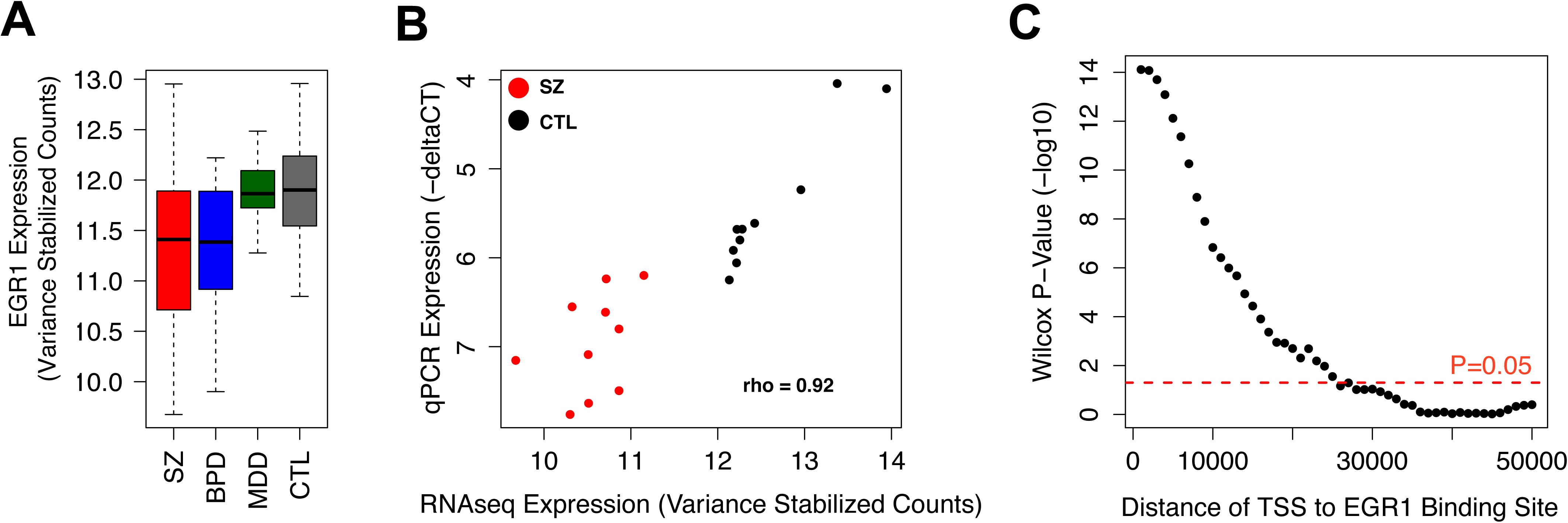
(A) Boxplots indicating relative expression of *EGR1* in the AnCg of SZ (red), BPD (blue), MDD (green), and CTL (gray). (B) Correlation plot comparing RNA-seq measured expression level of *EGR1* to qPCR measured expression in 10 SZ (red) and 10 CTL (black) patients. (C) Wilcox p-values resulting from comparing the degree of differential expression (based on DESeq2 p-values) of genes whose TSS are within the indicated distance to an *EGR1* binding sites compared to to genes whose TSSs are greater than the indicated threshold.

### Cell type specific changes

In addition to dysregulation of broadly acting transcription factors, another mechanism that can drive large-scale transcriptional changes in bulk tissue is alterations in constituent cell type proportions. Previous studies have observed decreases in neuron density and increased glial scarring in psychiatric disorders [36,37]. To test for signs of changing cell populations in our dataset we applied a method to deconvolute RNA expression data and estimate cell type proportions. Darmanis et al. identified genes capable of classifying cells into the major neuronal, glial, and vascular cell-types in the brain based on single cell RNA sequencing. We used these gene sets to generate cell type indices using the median of normalized counts for each cell type-specific gene set. We tested these indices on purified brain cell populations (Zhang et al.) and *in silico* mixed cell populations to demonstrate their accuracy and specificity [28,29] (Figure S4).

Application of these cell type indices to patient AnCg expression data revealed a significant decrease in neuron specific gene expression (Wilcox p-value<0.05) and a significant increase in astrocyte specific expression (Wilcox p-value<0.05) in SZ and BPD patients compared to controls (Figures 3A-B). Other brain cell-type indices were not significantly different between psychiatric patients and controls (Figure S5). An alternate algorithm for cell type deconvolution, DeconRNASeq, showed similar results (Figure S6A,B).

Additionally, we showed that neuron-specific genes identified by Darmanis et al. are enriched for decreased expression in SZ compared to controls and astrocyte-specific genes are enriched for increased expression (Figure S6C). Again, these enrichments are specific to this gene set and are not reproduced by 1000 expression matched, randomly sampled gene sets (Figure S6D,E). Further supporting a decrease in neuronal gene expression, we found a significant negative correlation between gene expression changes in patient brains relative to control brains and the degree of neuron specific transcription (fold enrichment of neuronal gene expression over other cell types) (SZ Rho=-0.50 and BPD Rho=-0.41, p-value<1E-16, SZ shown in Figure 3C).

**Figure 3.**
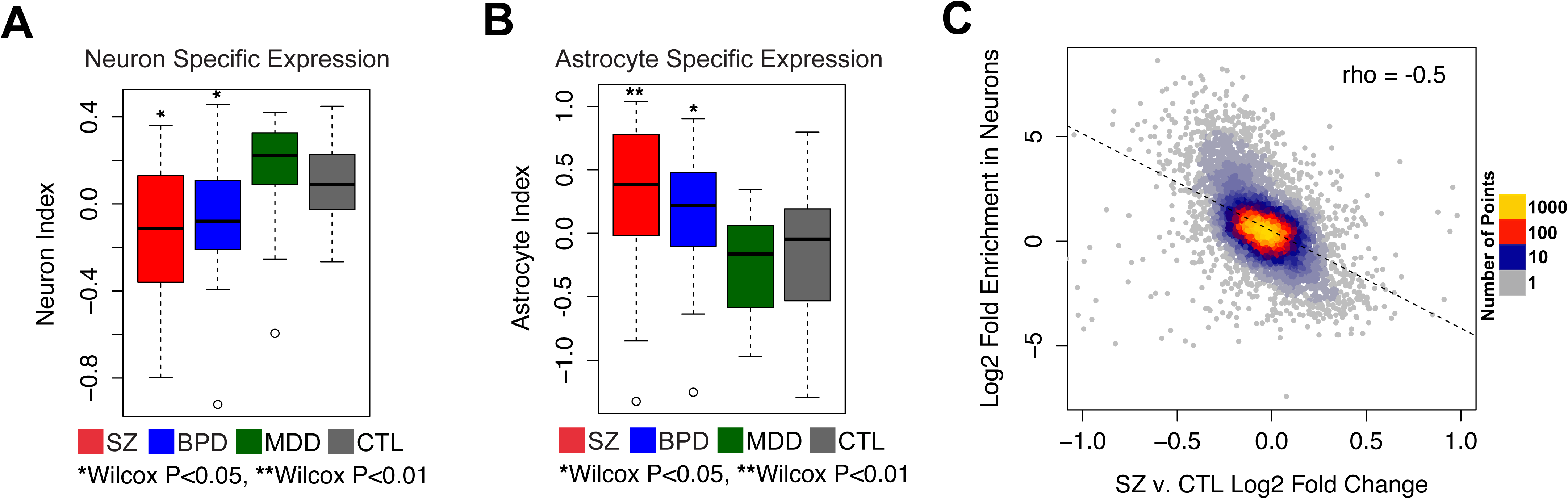
Boxplots indicating z-scored neuron-(A) and astrocyte-(B) specific expression indices in the AnCg for SZ (red), BPD (blue), MDD (green), and CTL (gray) individuals. (C) Correlation plot comparing the log2 expression fold change between SZ and CTL patients in the AnCg (X-axis) and the log2 fold change in gene expression from dissected neuron populations compared to all other dissected brain cell types (astrocytes, oligodendrocytes, endothelial cells, and microglia) for each transcript measured by Zhang et al.

### Transcriptomic changes reflected in altered metabolomic profiles

To assess the biochemical consequences of expression changes, we used 2D-GCMS to measure metabolite levels in 86 of the AnCg samples (sufficient tissue was unavailable for 10 samples). We measured and identified 141 unique metabolites (Table S8). We found no metabolites reached statistical significance (FDR<0.05), however 8 metabolites had an FDR<0.1 when comparing SZ to CTL. Similar to our gene expression analysis, metabolite levels (Table S9) successfully differentiated SZ and BPD patients from CTLs (Figure 4A), while MDD metabolite profiles were very similar to CTLs. Several of the most significant metabolites, including GABA, are known to be relevant to BPD and SZ (Figure 4B) [38]. Furthermore, GABA/glutamate metabolite ratios correlate strongly with average *GAD1* and *GAD2* expression levels measured by RNA-seq (Rho = 0.413, p-value=0.007, Figures 4C-D). This metabolite-gene relationship is consistent with previous multi-level phenomic analyses [39] and demonstrates realized biochemical consequences from altered gene expression. Notably, reductions in GABA could coincide with loss of neuron specific gene expression as suggested by the RNA-seq data. Integrated pathway analyses of metabolite and gene expression data revealed disruption of synaptic and neurotransmitter signaling in SZ compared to CTL (Figure S7, Table S10).

**Figure 4.**
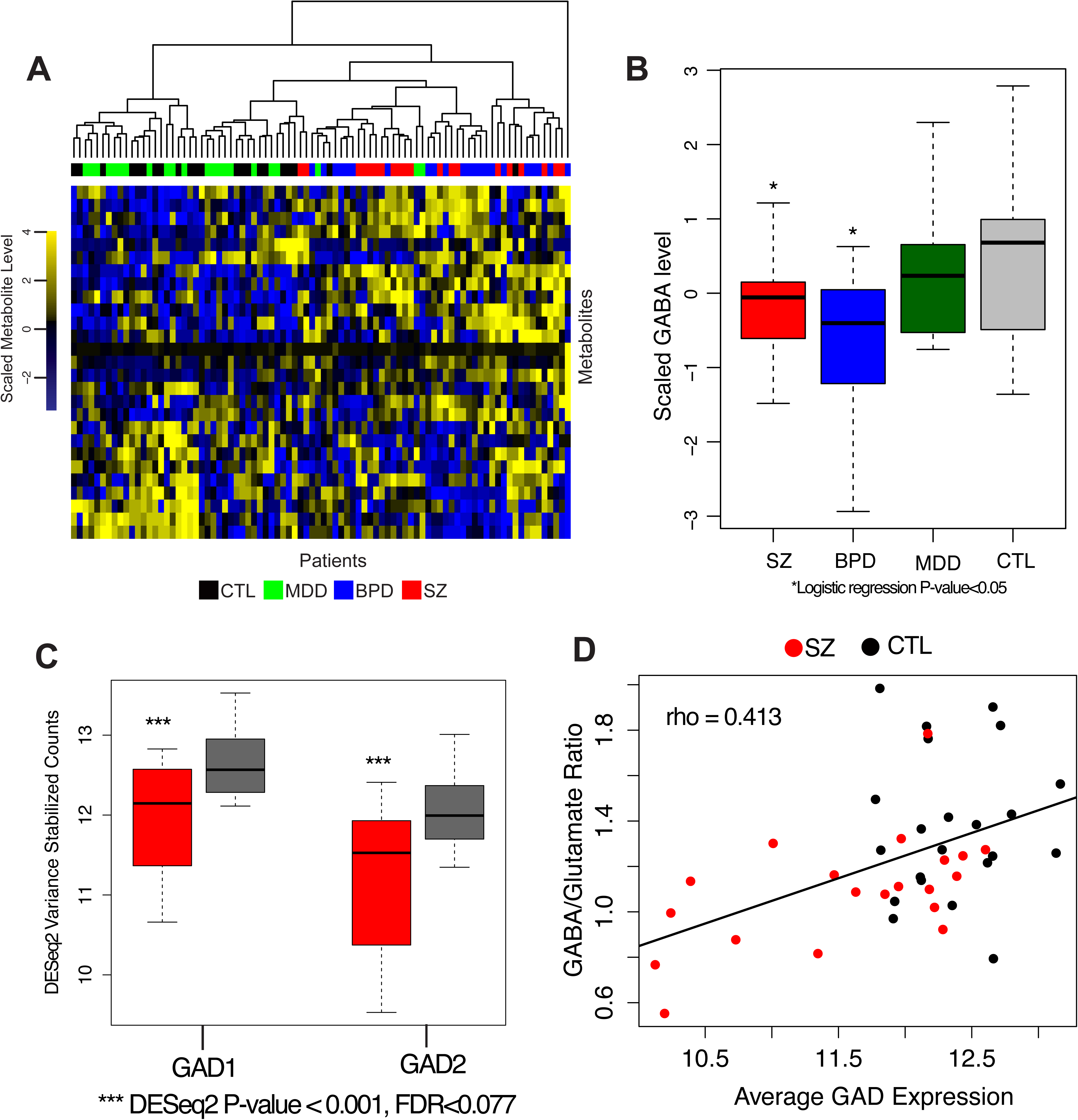
Hierarchical clustering of SZ (red), BPD (blue), MDD (green), and CTL (black) individuals using the top ten most significant metabolites for each case-control comparison (for a total of 30 metabolites). (B) Boxplots indicating z-scored GABA metabolite levels. (C) Boxplots indicating relative expression of GAD1 and GAD2 enzymes in the AnCg of SZ (red) and CTL (gray) patients. (D) Correlation plot comparing average GAD1 and GAD2 expression and the GABA/Glutamate metabolite level ratio in the AnCg of SZ (red) and CTL (black) individuals.

## Discussion

Here, we describe a large transcriptomic dataset across three brain regions (DLPFC, AnCg, and nAcc) in SZ, BPD, and MDD patients, as well as CTL samples matched for agonal state and brain pH. In MDD, we do not identify any genes that meet genome-wide significance for differential expression between cases and controls in any brain region. This finding agrees with previous post-mortem RNA-seq studies [40], however sample size and the choice of brain regions examined likely contributed to our inability to replicate results from previous non-transcriptome wide sequencing based approaches comparing MDD to CTL in post-mortem brain [41]. One limitation of our study is that females are underrepresented at a rate of about 5:1. This reflects the increased chance of accidental death among males [42], but limits us in our ability to make more general conclusions about these disorders and to address known differences between the sexes as they relate to these disorders. We also do not have information on the smoking status for our cohort, which is an important covariate as smoking rates are higher among patients with psychiatric disorders and smoking has been demonstrated to effect gene expression [43,44]. Another potential limitation inherent to post-mortem cohort analyses is accounting for patient drug use. As detailed in Table S2, patient toxicology reports were positive for several prescribed and illicit drugs that were not present in CTL samples. As this is a bias inherent to psychiatric patients it is impossible to disentangle from non-treatment related disease patterns in a post-mortem analysis.

Another important limitation of post-mortem RNA-sequencing studies is RNA quality. We found a significant proportion of variation in our data to be associated with multiple alignment quality metrics. Significant effort went into controlling for potential sources of bias due to differences in RNA quality. We only included tissue from patients with an agonal score of 0 and who had a brain pH of 6.5 or greater. We also controlled for brain pH, post-mortem interval, and alignment quality in all differential expression analyses. Our study, as well as future post-mortem studies, could be improved by directly measuring RNA quality at the time of sample preparation (e.g. RNA integrity number (RIN)). Even with these caveats, we believe our data yield new insights contributing to a growing understanding of these disorders.

The most dramatic gene expression signals we observed were brain region-specific. The majority of disease-associated expression differences were seen in the AnCg of SZ compared to CTL individuals. The AnCg has been associated with multiple disease-relevant functions, including cognition, error detection, conflict resolution, motivation, and modulation of emotion [45–47]. We observed a striking overlap in SZ- and BPD-associated expression changes consistent with previous findings [38,48].

One of the more intriguing genes significantly down regulated (FDR<0.05) in both cohorts of SZ patients was the zinc finger transcription factor, *EGR1*. We provide evidence that this factor binds upstream of a genes with altered expression in SZ and are associated with decreased expression in SZ patients. Down regulation of *EGR1* has been previously described in the prefrontal cortex of post-mortem brain samples from SZ patients [49,50]. *EGR1* has also previously been associated with several phenotypes relevant to psychiatric disorder including neural differentiation [51], emotional memory formation [52], response to antipsychotics [53], and has recently been described as part of a transcription factor-miRNA co-regulatory network capable of acting as a biomarker in peripheral blood cells (PBCs) for SZ [54]. In mice, loss of *EGR1* has linked to neuronal loss in a model of Alzheimer’s Disease [55]. *EGR1* is also important for regulation of the NMDA Receptor pathway, which is critical for synaptic plasticity and memory formation and has been implicated in SZ in humans [56]. We believe a more detailed examination of genome-wide *EGR1* occupancy in post-mortem brain tissue or cultured neurons could yield additional information and assessment of the functional consequences of *EGR1* perturbation is required to confirm this factor’s role in SZ pathogenesis.

We also see evidence for depletion of neuron-specific genes and increased levels of astrocyte-specific genes in SZ and BPD patients. This observation is further supported by metabolomic analysis of the AnCg, which found a concordant decrease in GABA levels in BPD and SZ individuals. Neuronal depletion has been previously described in SZ [36,37]. Insufficient tissue remains from our patient cohort to validate computational cell type predictions immunohistochemically, however our data strongly suggests that future post-mortem studies should be cognizant of cell type heterogeneity across patient samples. The method for cell type composition estimation is limited in its accuracy to estimating only the major classes of cells present. Genes represented in cell types present at only a small minority could be over or under-represented using this technique. Based on these results, future studies should consider using robust techniques for assessing tissue composition to examine potential cell type proportion differences between disease cohorts and to identify which transcriptional changes occur in conjunction with, and independent of, those differences.

We observed very few or no significant expression differences in the DLPFC and nAcc, which contradicts several previous studies [57,58]. We do not intend to claim that no transcriptional changes occur in these brain regions as our study was designed to broadly compare transcriptional alterations across multiple brain regions in multiple psychiatric disorders, thereby sacrificing exceptional sample sizes in any single disorder in any specific brain region. However, our data does suggest that of the regions we tested, the strongest transcriptional changes occur in the AnCg of SZ patients. Moreover, this data provides a useful resource for future studies facilitating the testing of preliminary hypotheses or validation of significant findings.

## Conclusions

Our study provides several meaningful and novel contributions to the understanding of psychiatric disease. We provide a well-annotated data set that has the potential to act as a broadly applicable resource to investigators interested in molecular changes in multiple psychiatric disorders across multiple brain regions. We have conducted an extensive characterization of the molecular overlap between SZ and BPD at the gene expression and metabolite level across multiple brain regions. We provide a high confidence set of genes differentially expressed between SZ and CTL individuals utilizing two independent cohorts and highlight down regulation of *EGR1* as a potential driver of broader scale transcription changes. We also establish that a significant proportion of transcriptome variation within SZ and BPD cohorts is correlated with expression changes in previously identified cell type-specific genes.

## List of abbreviations

RNA-seq: RNA sequencing
GABA: gamma-Aminobutyric acid
GWAS: genome-wide association study
SZ: schizophrenia
BPD: bipolar disorder
MDD: major depression disorder
CTL: control
AnCg: anterior cingulate gyrus
DLPFC: dorsolateral prefrontal cortex
nAcc: nucleus accumbens
GO: gene ontology
ChIP-seq: chromatin immunoprecipitation with DNA sequencing
PCA: principal component analysis

## Declarations

### Ethics approval and consent to participate

Sample collection, including human subject recruitment and characterization, was conducted as part of the Brain Donor Program at the University of California, Irvine, Department of Psychiatry and Human Behavior (Pritzker Neuropsychiatric Disorders Research Consortium) under IRB approval (UCI 88-041, UCI 97-74).

### Consent for publication

Not applicable

### Availability of data and materials

The datasets supporting the conclusions of this article are available in the GEO repository, GSE80655.

### Author’s Contributions

HA, SJW, AFS, WEB, JDB, HK, SJC and RMM conceived of study

KMB, RCR, BNL, SJC, AAH, MH, JZL and RMM designed the experiments

EGJ performed brain dissections

PMC procured the brain tissue samples

MPV analyzed pH on all cases and matched the 4 cohorts

DWM obtained demographic and clinical data on all subjects through analyses of medical records and next-of-kin interviews

NSD, JG, and KMB collected RNAs and performed Tn-RNA-seq library construction

RCR and BNL analyzed the RNA-seq data

RCR and SJC performed and analyzed metabolomics experiments

KMB, RCR, and BNL wrote the first draft of the paper

JZL, BGB, WEB, SJW, SJC, HA and RMM contributed to the writing of the paper

All authors read and approved the final manuscript.

## Acknowledgements

We thank Marie Kirby, Brian Roberts, Mark Mackiewicz, and Greg Cooper for many helpful discussions and comments on the manuscript, and all the members of the Pritzker Neuropsychiatric Disorders Consortium for their support and advice.

## Supplementary Figure Legends

**Figure S1.** A) Principal components analysis of all 281 brain tissues. AnCg (red squares), DLPFC (blue triangles), nAcc (green circles). B) Principal components analysis of all 281 brain tissues. CTL (gray squares), BPD (blue triangles), MDD (green circles), SZ (red triangles). (C) Principal components analysis of all 281 brain tissues after correcting RNA-seq data for alignment quality. CTL (gray squares), BPD (blue triangles), MDD (green circles), SZ (red triangles).

**Figure S2.** Principal components analysis of all AnCg (A,D), DLPFC (B,E), and nAcc (C,F) samples before (A-C) and after (D-F) correction for RNA-seq alignment quality. (G-I) PC1 values in CTL (gray), BPD (blue), MDD (green), and SZ (red) patients pre- and post-RNA-seq alignment quality correction in the AnCg (G), DLPFC (H), and nAcc (I).

**Figure S3.** GO-term analysis for genes differentially expressed in SZ vs. CTL in AnCg (FDR<0.05). Up-regulation (red circles), down-regulation (blue circles).

**Figure S4.** Using cell-type specific index based on single cell sequencing, we calculated the index for purified populations of (A) neurons, (B) astrocytes, (C) oligodendrocytes, (D) microglia, and (E) endothelial cells. In all cases the calculated indexes were specific to the purified population (F) Neuron and astrocyte indices were used to predict the proportion of each cell type in *in silico* mixed cell-type populations. (G) Compared to the Darmanis et al. indexes, 10,000 randomly generated gene sets do not predict cell type proportions. Mean values with standard deviation are plotted (H, I) Histogram of mean squared error of null index cell type proportion predictions for mixed neuron and astrocyte transcriptomes with Darmanis et al. gene performance indicated in red. The Darmanis gene set far outperforms any randomly generated gene sets.

**Figure S5.** Boxplots of endothelial (A), microglia (B), and oligodendrocyte (C) cell type indices in SZ (red), BPD (blue), MDD (green), and CTL (gray) individuals using indices derived from Darmanis et al.

**Figure S6.** (A,B) Estimated neuron (A) and astrocyte (B) cell type proportions using the deconRNAseq deconvolution algorithm in SZ, BPD, MDD, and CTL individuals. (C) Volcano plot of cell type specific transcripts in the Darmanis et al. gene sets show altered expression in SZ v CTL. Neuronal transcripts are enriched for loss of expression and astrocyte-specific transcripts are enriched for increased expression (D,E) Histograms representing the distribution of median log2 fold change for expression level matched gene sets to neuron (D) and astrocyte (C) specific genes. Red lines indicate median log2 fold change observed for neuron- and astrocyte-specific genes respectively.

**Figure S7.** Integrated KEGG pathway analysis of metabolite and RNA-seq differences between SZ and CTL patients. Top 10 pathways are shown for metabolite, gene and combined analysis.

